# Gibbs Free Energy of Protein-Protein Interactions Correlates with ATP Production in Cancer Cells

**DOI:** 10.1101/498121

**Authors:** Stefan M. Golas, Amber N. Nguyen, Edward A. Rietman, Jack A. Tuszynski

## Abstract

In this paper we analyze several cancer cell types from two seemingly independent angles: (a) the over-expression of various proteins participating in protein-protein interaction networks and (b) a metabolic shift from oxidative phosphorylation to glycolysis. We use large data sets to obtain a thermodynamic measure of the protein-protein interaction network, namely the associated Gibbs free energy. We find a reasonably strong inverse correlation between the percentage of energy production via oxidative phosphorylation and the Gibbs free energy of the protein networks. The latter is a measure of functional dysregulation within the cell. Our findings corroborate earlier indications that signaling pathway upregulation in cancer cells are linked to the metabolic shift known as the Warburg effect, hence these two seemingly independent characteristics of cancer phenotype may be interconnected.

## Introduction

Continued growth and proliferation under hypoxic conditions in cancer cells entails significant energy and material demands. Since the publication of Warburg’s foundational paper on cancer metabolism in 1925, much research has focused on the role of glycolysis as the primary source of energy in cancer cells. However, recent reviews (Jose et al. 2011; Zu and Guppy, 2004; Moreno-Sanchez et al. 2007; Vander Heiden and DeBerardinis, 2017) and meta-analyses of studies on cancer metabolism have shown that many cancers rely heavily on ATP produced from oxidative phosphorylation (OXPHOS). These studies identify a wide range of experiments, including ^13^C and ^14^C labeled fuel sources (e.g. Marin-Valencia, et al., 2012) to corroborate their conclusions. The work of Moreno-Sanchz et al. (2009, 2007), and Rodriguez-Enriquez et al. (2000), which demonstrate high levels of oxygen consumption in tumors, as well as point out flawed methodologies in several papers seeming to validate the Warburg effect. Zu and Guppy (2004), in particular, emphasize that ATP generation should be tracked from O_2_ consumption and lactate production at a minimum, and that studies which fail to track either of these molecular species are inherently flawed, in their perspective, on the roles of glycolysis versus oxidative phosphorylation. They further argue that cancer metabolism is glycolytic only insofar as the environment is hypoxic. Additionally, Wu et al. (2016) note that lactate buildup causes a shift back to an oxidative phosphorylation phenotype in numerous cancer cell lines. It is clear that beyond merely a shift from oxidative phosphorylation to glycolytic metabolism, ATP generation in cancer is highly dependent on cell lineage and environmental constraints.

The limitless and self-sufficient proliferative potential exhibited by cancer cells entails significant metabolic demands. Many cell activities carried out during mitosis are very energy-intensive and require considerable stores of ATP, particularly the formation and disassembly of the mitotic spindle and unfolding and refolding of the DNA double helix. The sheer quantity of mass required for cell replication also involves considerable material demands, thus angiogenesis is frequently observed in cancer tumor environments in order to provide additional nutrient supply through the recruitment of blood vessels. The persistently high rates of energy and material consumption in tumors results in fierce competition for resources with surrounding host cells, a driving force behind cancer’s invasiveness and eventual lethality. Tumors that reproduce most prolifically have optimized metabolic pathways that seize substrates in their microenvironment and hence pose the greatest danger to the host (Guppy et al, 2002).

Jose et al. (2011) argue that it is time for a unifying theory describing ATP production in cancer cells. This, more comprehensive theory, would include bioenergetic profile of patients’ tumors and thus enable a more personalized treatment of tumors based on specific energetic and biochemical pathways (Gatenby and Gillies, 2007; Rietman, et al. 2017). In this manuscript take a step in the direction of a quantitative understanding of cancer metabolism and calculate a thermodynamic measure known as Gibbs free energy for a number of cancer cell types using available literature data. We then show that the Gibbs free energy of the protein-protein interaction networks for these cancer cell types is reasonably closely correlated (R = -0.716) with the percentage of ATP contribution obtained from OXPHOS. This provides a mechanistic interpretation of the metabolic energy demands correlating with protein expression data. High percentage of OXPHOS contribution is associated with relatively low level of malignancy while high values of the over-expressed protein-protein interaction network’s Gibbs free energy indicate the opposite, hence an inverse correlation relationship. The protein-protein interaction network complexity as measured by degree entropy has earlier been shown to correlate to a similar degree with cancer lethality (Breitkreutz et al. 2012). Putting this together links protein expression dysregulation to increased metabolic energy demands with a glycolytic shift as its characteristic and eventually to poor survival prognosis.

## Methods

Gibbs free energy, *G*, is one of the fundamental thermodynamic functions of state, which describe the state of a macroscopic molecular system independent of the path or process that has taken the system to this state. This thermodynamic quantity is especially relevant in the context of systems kept at a constant temperature and able to exchange both energy and molecules with the surrounding environment. Gibbs free energy is most readily described in terms of chemical potentials defined for each molecular species represented in the system. If one protein, A, is in high abundance and another protein, B, needed for a specific biochemical reaction involving that first protein is in low abundance, then there is said to be a chemical potential difference between A and B. For a system of interacting proteins, we may take the reference state where *G* = 0 to be the condition where no interactions are occurring. If we have a system of fixed volume, then *G*_*i*_ = *c*_*i*_*Ḡ*_*i*_ where *c*_*i*_ is the molar concentration of protein *i* and *Ḡ*_*i*_ is the Gibbs energy per mole of protein. Here, *i* refers to a set of individual proteins {*i_*1*_, i_2_ … i_N_*} within the protein-protein interaction network.

The values of Gibbs free energy for a cell can be calculated from mRNA expression concentrations, a surrogate for protein concentrations, and overlaid on an expression network (Rietman, et al., 2017). The Gibbs values used in this paper were calculated using protein-protein interactions from BioGRID, a curated database (https://thebiogrid.org/). Gene expression data for several cancers were obtained from TCGA (https://cancergenome.nih.gov/). The cancers we selected from TCGA spanned a range of cancer-stage from I to IV. These mRNA expression data were used as a surrogate for protein concentrations. **Eq. 1** below is the formula for obtaining the Gibbs energy for a given protein *i*, where *c_i_* refers to the normalized concentration of protein *i* estimated from its gene expression level, and *C_j_* refers to the concentration of the *j*th interaction partner. Normalization was performed by dividing the concentration of each gene transcript by the highest concentration in that sample, such that values of *c_i_* range from 0 ≤ *c_i_* ≤ 1. Using this formula, the Gibbs free energy of a protein will be some *G_i_*≤ 0, the magnitude of which is associated with the amount of that protein bound to interaction partners. **Eq. 2** below is the formula for obtaining the Gibbs energy of the whole network. These formulae were implemented in a Python 2.7 script running NetworkX, SciPy, and NumPy packages.

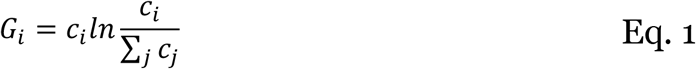

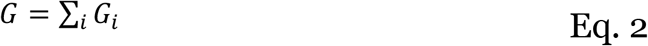

## Results and Discussion

As described in the **Methods** section we used data from Rodriguez-Enriquez et al (2000) for OXPHOS - ATP production, and TCGA transcription data (https://cancerqenome.nih.gov/) for our analysis of the corresponding Gibbs free energy values. **Figure 1** shows a negative correlation (R=-0.716) between Gibbs free energy and the percentage of ATP derived from oxidative phosphorylation. The graph suggests that certain cancers obtain their ATP predominately from OXPHOS and others from glycolysis.

**Figure 1.**
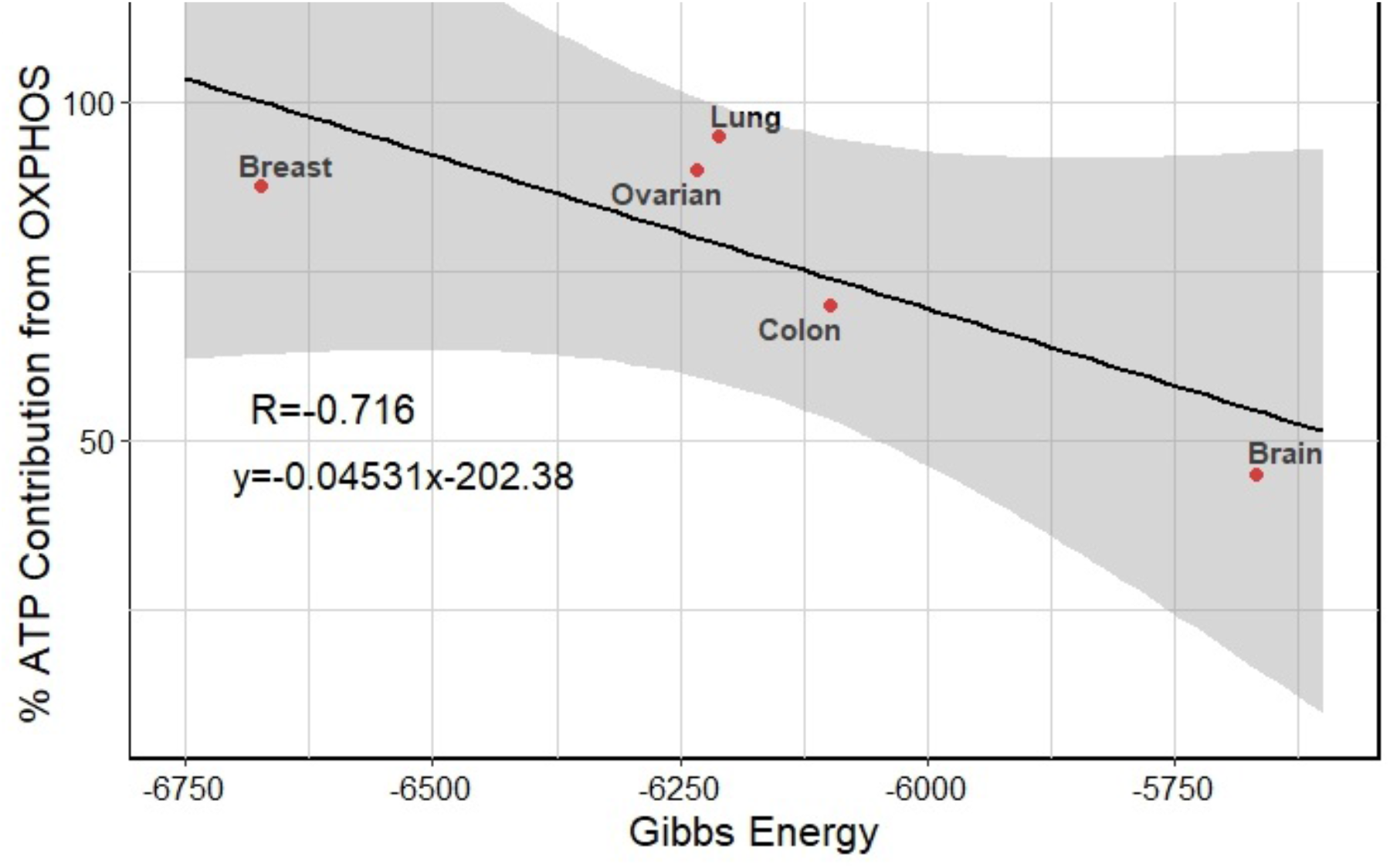
Cancer metabolism by type. Y-axis data refer to the overall contribution to ATP generation from oxidative phosphorylation and were determined from a number of metabolic flux analyses whose results are tabulated in Rodriguez-Enriquez et al (2000). The percentages may be contrasted with % ATP from glycolysis. “Lung” refers to lung squamous cell adenoid cancer, and “brain” refers to glioblastoma.

The extended metabolic pathway used in OXPHOS, and the high number of subunits found in the enzymes of this pathway, are likely to be significant contributing factors to the observed results in Figure 1. The networks (from KEGG, https://www.genome.jp/kegg/) for the OXPHOS machine are shown in Figure 2a. By contrast the glycolysis network is shown in Figure 2b. The difference between the highly symmetric OXPHOS network geometry and the much less regular glycolytic network architecture is striking.

**Figure 2.**
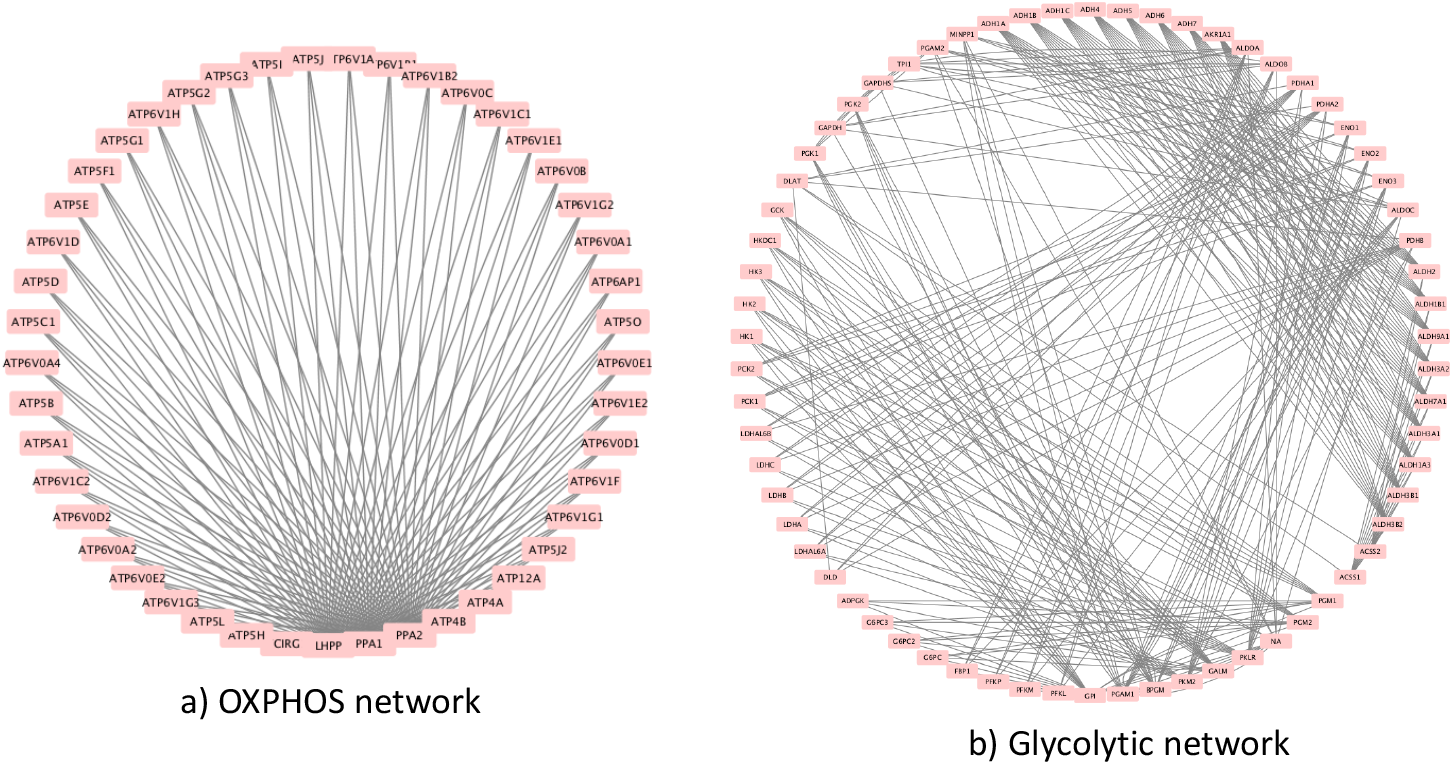
KEGG networks converted to protein-protein interaction networks with KEGGgraph and plotted in Cytoscape.

In the following we describe several network statistical measures; details of their algorithms can be found in Newman (2010). The first thing to notice about the two networks is the large difference in connectivity. The OXPHOS network has three key nodes, LHPP, PPA1 and PPA2. These three proteins connect to 42 other proteins to form large molecular machine parts and then these three machines interact for the ATP production. In contrast, the glycolytic network has one hub, GPI, connected to 23 other proteins in the network. The shortest path in the OXPHOS network is 1.05 for the three hub nodes and the remaining nodes have a shortest path of 1.9. On the other hand, the shortest path in the glycolytic network is 2.8 and increasing to 5.8. The clustering coefficient is 0.0 for the OXPHOS and 0.66 for the glycolytic network. Lastly, the betweenness centrality is found to be 0.303 for the three hub nodes in the OXPHOS network and <10^-5^ for the remaining nodes. In the glycolytic network the betweenness centrality is in the range from 0.46 to 0.0017 (largest to smallest). These network measures imply that there is greater free energy utilized in the OXYPHOS network relative to the glycolytic network.

The high number of subunits in OXPHOS enzymes speaks to the intricacy of their operations relative to glycolysis; while glycolysis generates ATP from the group transfer potential of phosphate groups on substrate, OXPHOS complexes use electron tunneling and proton pumps to establish an H^+^ concentration gradient across the mitochondrial membrane. The chemical potential of each individual participant in these complexes may be higher than those in glycolytic monomers and dimers due to their coexpression alongside many interaction partners.

## Conclusion

According to the Warburg hypothesis, the aggressiveness of a tumor derives from the shift of metabolic energy production from OXPHOS to glycolysis. This shift is considered to be due to the competition between cells preferentially utilizing the glycolytic mode, and cells adopting the OXPHOS mode of energy production. In OXPHOS, the coupling between the redox reactions taking place in the mitochondria and ADP phosphorylation is electrical through the electron transfer mechanism in the mitochondrial wall protein complexes. Hence the metabolic rate is determined by bio-energetic parameters such as the proton conductance and proton potential of the metabolic network. In glycolysis on the other hand, ADP phosphorylation is linked chemically to individual enzyme-catalyzed reactions, so the rate of ATP production is a function of the flux through the pathway as a whole.

However, it is well known that ATP generation through glycolysis is less efficient than through mitochondrial OXPHOS processes. Hence a long-standing paradox is how cancer cells with their metabolic disadvantage can outcompete normal cells. In terms of biochemical reactions (Pelicano et al. 2006), mitochondrial respiration defects activate the Akt survival pathway through a mechanism mediated by NADH. Respiration-deficient cells harboring mitochondrial defects exhibit dependency on glycolysis, increased NADH, and activation of Akt, leading to survival advantage (Pelicano et al. 2006). This inter-relation between metabolic and signaling pathways in cancer cells has been indirectly shown in this paper via an inverse correlation between the percentage of OXPHOS utilization and the Gibbs free energy of the protein-protein interaction network.

Gibbs free energy is a topological measure on protein interaction networks, which remains to be fully explored in its usefulness for identifying targets for cancer treatment. By targeting proteins based on their participation in multi-subunit complexes, it may be possible to diminish the functionality of entire complexes or pathways using a single customized drug. Drugs targeting proteins with many highly expressed interaction partners should affect the functionality of these partners by proxy, producing a cascade of diminished functionality that may significantly increase the likelihood of cell death.

## Acknowledgments

JAT gratefully acknowledges generous support from NSERC (Canada).

